# Extracellular matrices of bone marrow stroma regulate cell phenotype and contribute to distinct stromal niches *in vivo*

**DOI:** 10.1101/2023.01.19.524473

**Authors:** Andrew Stone, Emma Rand, Gabriel Thornes, Alasdair Kay, Amanda Barnes, Ian Hitchcock, Paul Genever

## Abstract

The heterogeneity of bone marrow stromal cells (BMSCs) has been revealed more in recent years through the advent of single cell RNA sequencing. However, protein level characterisation is likely to provide a deeper understanding of the functions of individual subsets and may reveal insights into the co-ordination of the cell phenotype maintaining niche.

Here, by analysing heterogeneity in BMSC populations using human stromal cell lines to model extremes of cell morphology and migration characteristics, we identified plastic cell phenotypes that can be modified through secreted proteins. Transfer of secreted signals from a differentiation-competent stem cell phenotype was able to stimulate migration in a slow-moving stromal cell, observed via label-free ptychography. Subsequent untargeted proteomic interrogation of the secreted factors from these cell lines identified a highly significant enrichment of extracellular matrix (ECM) protein production by the differentiation-competent cells compared to non-stem cells. The most highly enriched proteins, aggrecan and periostin, were identified on the endosteal surfaces of mouse and human bone, underlying CD271+ stromal cells in the latter, indicating that they may represent key non-cellular niche-components important for cell maintenance and phenotype. ECM from stem cells was further capable of enhancing migration in non-stem cells in a focal adhesion kinase-dependent manner.

Overall, we demonstrate the importance of the ECM in co-ordination of cellular phenotype and highlight how non-cellular components of the BMSC niche may provide insights into the role of BMSCs in health and disease.

## Introduction

The bone marrow microenvironment is complex, with interplay and heterotypic signalling between haematopoietic and non-haematopoietic compartments [1, 2]. The study of bone marrow stromal cells (BMSCs) within this environment has often focused on how BMSCs interact and communicate with other cell types, with particular attention to their role in skeletal homeostasis, haematopoietic control and immunomodulation [2-7].

We and others have previously reported considerable heterogeneity in stromal populations, in terms of morphological and functional characteristics [8, 9]. Work in both mice and humans has provided evidence for a carefully co-ordinated developmental tree of BMSCs that is critical to skeletal lineage differentiation and bone marrow architecture [10, 11]. Recent developments in single-cell profiling have facilitated the interrogation of stromal diversity, with several reports of complex, heterogeneous subsets [12-18]. However, there has been little work to investigate how these stromal subsets interact with one another to influence larger scale remodelling and inflammatory responses in healthy and disease states. BMSCs are capable of mediating both pro- and anti-inflammatory effects, and ample evidence suggests a correlation between cellular morphology and function [19-21].

We previously reported the development of a panel of human telomerase reverse transcriptase (hTERT) immortalised clonal BMSC lines that partially model stromal heterogeneity in bone marrow [22]. These include the Y201 line which exhibits classical stem cell-like tri-potent differentiation capacity, and the Y202 BMSC line, that is nullipotent and has pro-inflammatory characteristics. Both of these BMSC lines express cell surface proteins described by Dominici, et al. [23] well as the commonly reported marker leptin receptor (LEPR). However, Y201 and Y202 BMSCs display considerable variation in morphology, migration, transcriptional profiles and function, highlighting a need for further refinement of stromal identity [24].

Heterogeneous stromal cells are likely to reside in subtype-specific locations *in vivo* and their local environment will have considerable influence on cell phenotype. There is also significant interest in the role that BMSCs play in the haematopoietic niche, therefore defining the composition of specific niche environments would aid understanding of their function, in particular the contribution of non-cellular components such as cytokines and extracellular matrix (ECM). There is also specific relevance for understanding disease pathologies; for example, de Jong, et al. [25] showed evidence for involvement of different subsets of BMSCs in multiple myeloma.

Here we used our immortalised BMSC lines to examine phenotypic stability. We demonstrate that heterogeneous BMSC sub-populations are inherently plastic both in terms of cell morphology and migratory characteristics and that this plasticity is inducible through the exposure to secreted factors from different stromal subsets, in particular ECM components. Our findings add to our understanding of the mechanisms that determine the onset and resolution of heterogeneity in different cell and tissue contexts. Furthermore, we demonstrate differential contribution of BMSC subsets to ECM production *in vitro* and highlight candidate components of a putative stem cell-supporting niche *in vivo*, which will prove important for understanding disease development, identification of functional subpopulations and for production of *ex vivo* expanded cells for therapeutic applications.

## Results

### Heterogeneous BMSCs have distinct morphologies and migratory characteristics

Through colony-forming unit fibroblastic (CFU-F) and related assays our group and others have identified morphologically distinct BMSC subtypes in primary donor populations. The morphology of colonies and of individual cells within colonies could reflect and/or be predictive of biological function. The immortalised BMSC lines, Y201 and Y202, have different cellular morphologies; Y201 cells have a typical elongated, bipolar stromal morphology, whereas Y202 cells are round, flat and spread (Fig.1A). Using the program CellProfiler, we quantified aspects of cellular morphology from label-free ptychographic images and revealed a significantly larger length:width ratio in Y201 versus Y202 BMSCs (3.59±0.072 vs 2.016±0.051, mean±SD, p<0.0001), whereas Y202 cells had an increased average cell area (p<0.0001) versus Y201 cells (Fig.1B&C). We also observed differences in migratory phenotype, visualised in rose plots generated by tracking individual cells (Figs.1D) with Y201 cells moving nearly twice as far and more quickly on average (p<0.0001) (Fig.1E&F).

To determine whether differences in cell morphology correlated with cytoskeletal variations, we fluorescently labelled focal adhesions (FA) and actin in Y201 and Y202 BMSCs (Fig.1G). Phalloidin staining revealed criss-crossing actin networks in Y202 cells whereas Y201 appeared to have more aligned actin fibres. Quantification of FA size revealed that FAs in Y202 cells displayed a significantly increased mean area of 1.572μm^2^ versus 1.164μm^2^ per adhesion in Y201 (Fig.1H). As well as increased area per-adhesion, Y202 also had significantly more adhesions on average (Fig.1I)

We confirmed the presence of Y201-like and Y202-like populations in CFU-F cultures of primary BMSCs by building an analysis pipeline that could distinguish and classify cells based upon morphological phenotypes. Using the CellProfiler pipeline we identified contrasting phenotypes within the same culture of primary cells, including cells with Y201-like fibroblastic morphologies and Y202-like, flattened and spread morphologies (Fig.1J). Image analysis of three separate primary cultures nominally identified 48.5% of cells in primary cultures as “Y201” and 24.7% as “Y202” (Fig.1K). The remaining 26.1% was designated as unclassified, having a morphology somewhere between the two defined populations. We conclude that morphologically and functionally-distinct cell subsets co-exist in BMSC populations.

**Figure 1.**
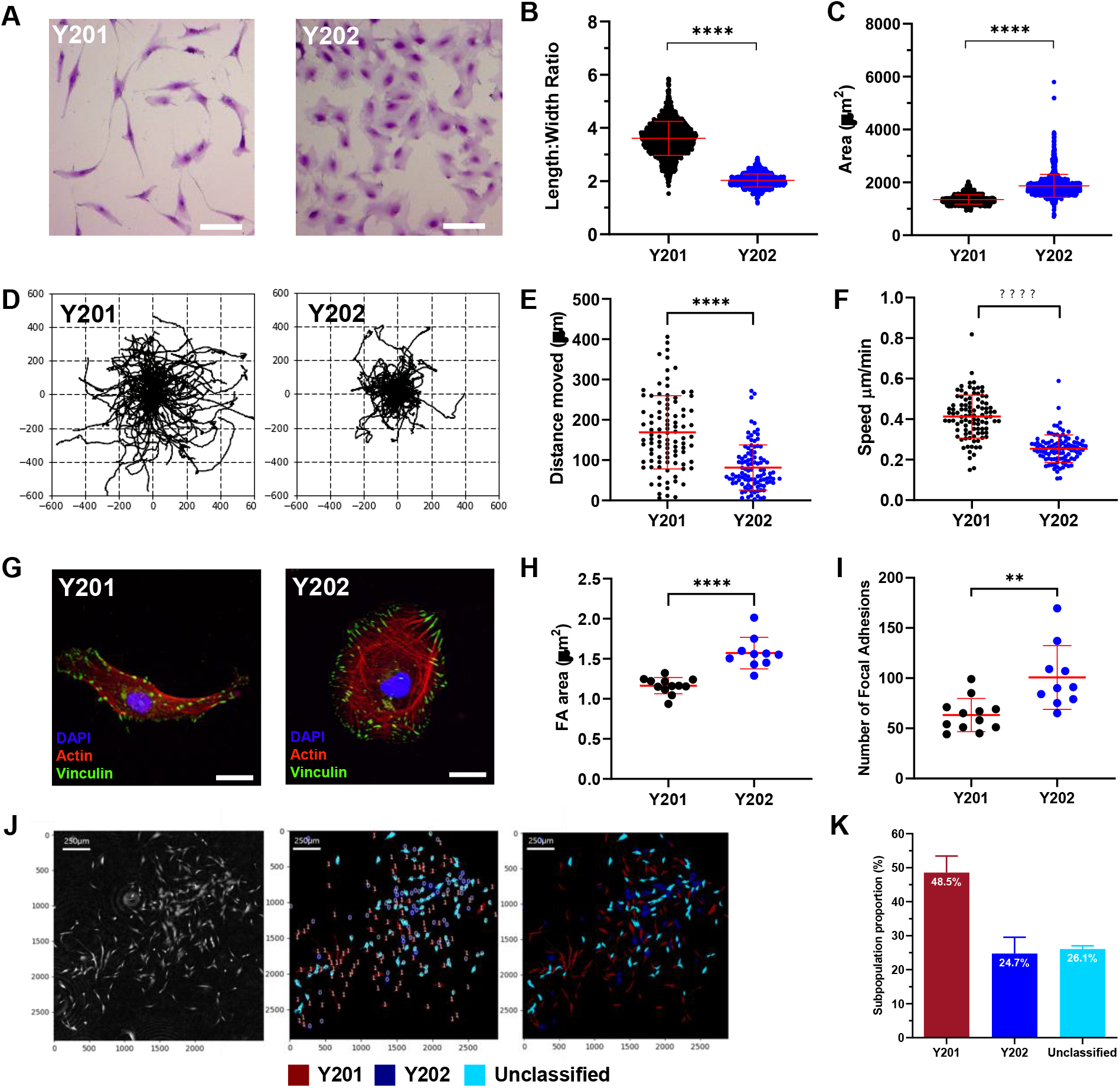
Image analysis of morphologies and migratory phenotypes in Y201 and Y202 BMSCs. A) Y201 and Y202 stromal cell subtypes stained with crystal violet and imaged by brightfield microscopy (scale bar = 50μm). B) Length:width ratios quantified from still frames from ptychographic images of Y201 and Y202 (T-test, P<0.0001, n=2418). C) Cell area quantified from still frames from ptychographic images of Y201 and Y202 (T-test, p<0.0001, n=2418). D) Rose-plots showing migratory profiles of Y201 and Y202 BMSCs. E), F) Quantification of migratory characteristics of Y201 and Y202 from ptychographic live-cell tracking. G) Representative immunofluorescence images of Y201 and Y202 cells showing focal adhesions (vinculin, green), actin (phalloidin, red) and nuclei (DAPI, blue), scale bar = 20μm H) Quantification of fluorescence images for mean focal adhesion (FA) area of Y201 vs Y202 (n=10-12) I) Number of focal adhesions per cell from Y201 and Y202 cells. J) Ptychography was used to build a CellProfiler pipeline that could classify primary cell populations based upon Y201 and Y202 morphological metrics. Representative images show phase-contrast images in the first frame which are overlayed to represent the classification of primary BMSCs. Red = Y201-like subtypes, blue = Y202-like subtypes, light-blue = unclassified. K) Quantification of Y201 and Y202-like subtypes identified in primary BMSC populations, all error bars = Mean±SD. *p≤0.05, **p<0.01, ***p<0.001, ****p<0.0001

### Secreted factors from Y201 BMSCs drive phenotypic switching in Y202 BMSCs

We hypothesised that the BMSC phenotype is plastic and at least in part regulated by the interactions of clonally-derived cell subtypes to determine the overall function of the population. We used the unique, quantifiable features of Y201 and Y202 cell lines as a model of BMSC heterogeneity to test this hypothesis. To determine the role of secreted factors on phenotype maintenance, we transferred conditioned media (CM) between Y201 and Y202, and monitored cell morphology and migration in CFU-F assays, focusing on the effects of the Y201 secretome on behavioural changes in atypical Y202 stromal cells. The varied morphology and inherent migration of some BMSC subtypes makes quantifying metrics from CFU-F assays complex. To overcome this, we developed a CellProfiler pipeline capable of accurately identifying colonies of various morphologies from scanned images of crystal violet stained CFU-F assays (Fig.S1). Exposure of Y202 cells to Y201-CM resulted in a significant increase in colony size compared to their own Y202-CM, and no conditioned media treated colonies (*p*=0.0157 and *p*=0.0018 respectively) (Fig. 2A&B). This increase in colony size appeared to arise from increased migration of cells resulting in colony spreading from the initiation point.

**Figure 2.**
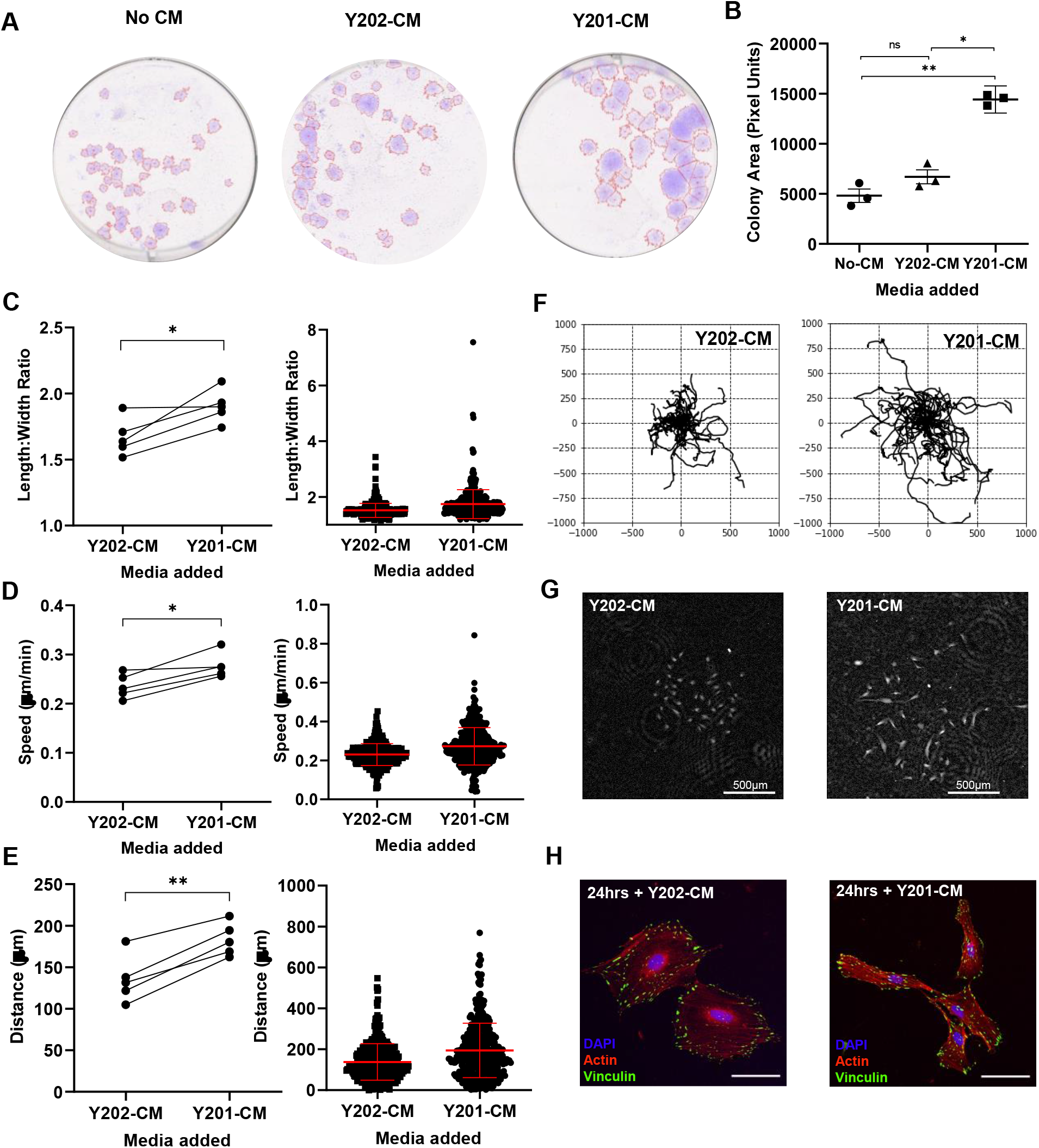
Effect of Y201 secreted factors on morphology and migration of Y202 cells. A) Representative images of crystal violet stained CFU-F cultures of Y202 grown in either unconditioned medium (no CM), Y201-conditioned medium or Y202-conditioned medium showing gross colony morphology B) Mean colony area of Y202 colonies treated with various conditioned media (ANOVA: *F* = 60.05, *d*.*f* = 1.12, 2.26, *p* = 0.0113) C) Mean length:width ratio from a single experiment (left) and multiple repeats (right) (n=5) D) Speed of mean cell movement from a single experiment and the mean speed from multiple repeats (n=5) E) Displacement distance of cells from tracking origin for a single experiment and the mean distance from multiple repeats (n=5). F) Rose-plots of cell migratory pathways after exposure to Y201 or Y202 conditioned media. G) Phase contrast image of typical colonies at assay endpoint H) Representative immunofluorescent images of Y202 cells treated with either Y201-CM or Y202-CM for 24 hours with focal adhesions (vinculin, green), actin (phalloidin, red) and nuclei (DAPI, blue), scale bar = 20μm. *p≤0.05, **p<0.01, ns = not significant. Error bars ± SEM

To quantify the effect of Y201-CM on Y202 cell migration further, we used a ptychographic imaging technique to track individual cells within colonies over time. From time-lapse imaging we observed that Y202 cells became more dispersed and elongated following exposure to Y201-CM compared with Y202-CM controls (Supplementary videos 1 and 2). In quantitative analyses we demonstrated that Y202 cells underwent morphological changes following exposure to Y201-CM, with a significant increase in length:width ratio (p=0.0293) (Fig.2C). Similarly, we found significant increases in migration speed (p=0.0141) and displacement distance (p=0.0012) for Y202 cells treated with Y201-CM versus Y202-CM (Fig.2D&E). Rose-plots from individually tracked cells illustrate the increased migration of Y202 cells with exposure to Y201-CM (Fig.2F) with examples of colonies at the assay endpoint shown in Figure 2G. By fluorescent staining we observed a change in morphology of Y202 cells to a more elongated Y201-like phenotype within 24 hours of exposure to Y201-CM (Fig. 2H), however, we saw no significant change in the size of focal adhesions between Y202-CM and Y201-CM treatments (p=0.46) (Fig.S2A). The mean focal adhesion size of Y202-CM (1.323μm^2^, n=16) and Y201-CM (1.395μm^2^, n=17) treated cells was notably between the sizes of the highly migratory Y201 and less migratory Y202 (1.572μm^2^ and 1.164μm^2^ respectively, shown in Fig.1H).

We repeated this CFU-F assay using *in vitro*-aged (>10 passages) primary BMSCs, which typically display reduced CFU-F activity compared to low-passage cells. We found that primary *in vitro* aged cells exposed to Y201-CM increased the number of colonies, albeit not significantly (*p*=0.0738), but the subsequent colonies grew significantly larger than unconditioned media controls (*p*=0.0133) while Y202-CM had no significant effect over standard culture conditions (Fig.S2 B&C). In one donor (K136), Y201-CM completely restored colony forming capacity.

### BMSC heterogeneity is reflected in variability of secreted factors

Having determined that exposure to Y201 secreted factors was able to drive changes in morphology and migration of Y202 cells, we interrogated the secretomes of these distinct cell subsets using LC-MS/MS. Remarkably, all 861 proteins we detected were expressed by both Y201 and Y202 BMSC subtypes. From this we identified 44 and 129 proteins with significantly increased expression in Y201 and Y202 BMSCs respectively (p<0.05) (Fig.3A). Using the cell region-based rendering and layout tool in cytoscape we confirmed the majority of our differentially expressed proteins have been annotated as appearing in the extracellular space (Fig.3B). The 44 proteins significantly elevated in Y201 versus Y202 are shown in Fig.3C ranked by normalised abundance. The most highly abundant proteins with significant fold changes were predominantly ECM components (e.g. FN1, COL6A1, BGN, DCN, THBS1), with notable elevated levels of periostin (POSTN) and aggrecan (ACAN) (71- and 104-fold higher than Y202 respectively, Fig. 3C arrows). Conversely, Lumican (LUM) was the most upregulated ECM component in Y202 cell secretome with levels 9.7-fold higher than Y201 cells (Fig.S3A). KEGG pathway enrichment of significantly upregulated proteins revealed strong correlation for Y201 secreted proteins in the ‘ECM-Receptor Interaction’ and ‘Focal Adhesion’ pathways while Y202 upregulated proteins demonstrated weak but significant correlation with the ‘Lysosome’ and various metabolic pathways (Fig.3D).

**Figure 3.**
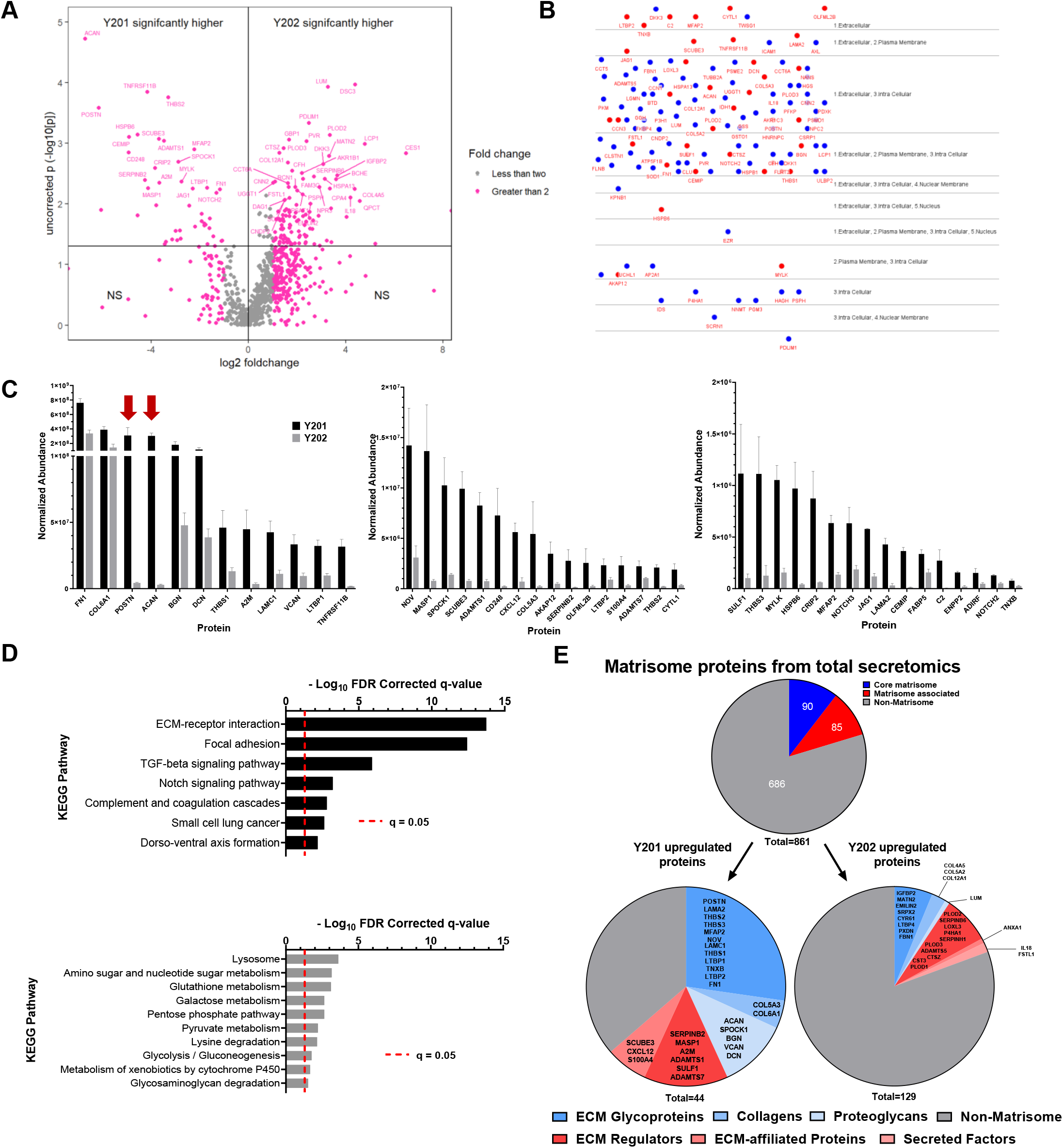
Analysis of Y201 and Y202 secretome composition by LC-MS/MS. A) Volcano plot showing proteins identified by LC-MS/MS in conditioned media from Y201 and Y202 cells. Proteins identified as significantly enriched by ANOVA n=3, p<0.05 are shown in upper quadrants. B) CEREBRAL layout of significantly differently expressed proteins from Y201 (red) and Y202 (blue) showing the majority have been annotated as found in the extracellular space and are likely secreted. C) Significantly enriched proteins secreted by Y201 versus Y202 represented in order of overall normalised abundance from LC-MS/MS. Graphs split for ease of interpretation while maintaining a linear scale, arrows indicate POSTN and ACAN, Means ± SEM. D) KEGG pathway analysis of significantly upregulated proteins in Y201 (top) and Y202 (bottom). E) All identified secreted proteins were annotated using the matrisome database (top) and categorized as “core matrisome” (blues), “matrisome-associated” (reds) and “non-matrisome” (grey). Significantly enriched proteins from secretomics for Y201 (left) and Y202 (right) are shown with matrisome annotations in the respective subcategories listed.

The recurring references to ECM in KEGG pathway enrichment was investigated further by comparing all proteins identified in LC-MS/MS of Y201 and Y202 secretome against the matrisome, a curated database of human proteins known to contribute to or associate with ECM through either structure, interaction, or regulation [26]. From 861 proteins identified in the secretome, 175 (20.3%) were annotated in the matrisome, with 85 labelled as “core matrisome” and 90 as “matrisome associated” (Fig.3E). Chi-squared tests revealed significant enrichment for matrisome proteins (28 observed versus 8.9 expected) in the Y201 secretome (χ2=50.97, d.f.=1, p<0.0001). Notably, Y202 significantly upregulated proteins did not differ significantly from expected amounts (χ2=0.1576, d.f.=1, p=0.69). Of the 175 matrisome proteins in the total secretome, 122 proteins did not differ significantly between Y201 and Y202 (Fig.S3B).

### Secreted ECM products from Y201 are found predominantly in stromal compartments *in vivo*

We used the ECM proteins identified in the secretomic screen as candidate markers of a Y201-like stromal cell niche. Using immunofluorescent labelling we identified expression of four ECM proteins differentially upregulated by Y201 BMSCs (collagen-VI, biglycan, aggrecan and periostin) in sections of mouse and human bone. All four ECM proteins were found lining trabecular bone, in addition networks of collagen-VI were also identified throughout the marrow (Fig. 4 A&B). Very similar distribution patterns were observed in mouse bone (Fig. S4).

**Figure 4.**
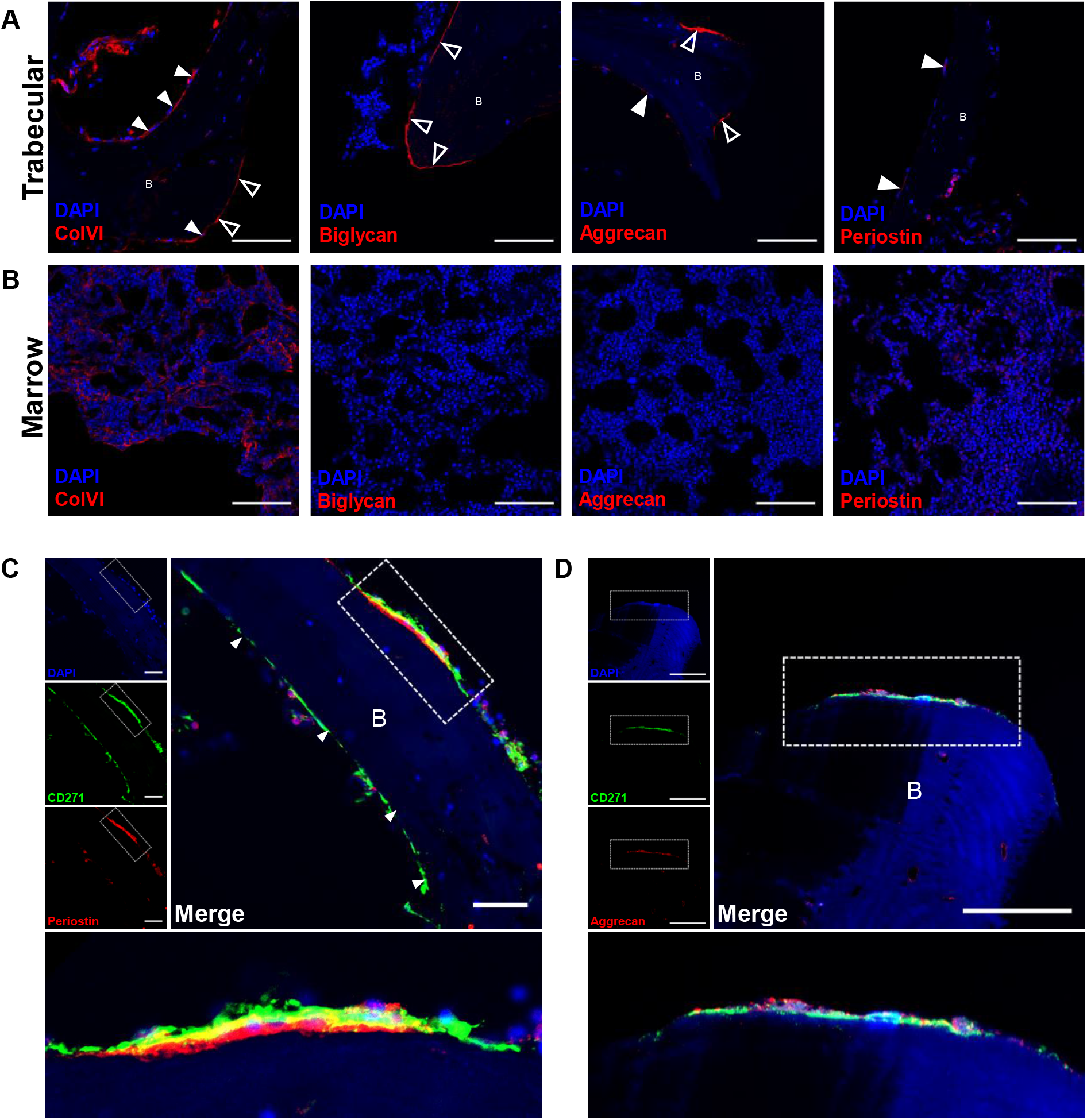
Immunofluorescent imaging of human bone marrow to identify Y201 BMSC-associated ECM proteins. A) regions of human trabecular bone with fluorescently labelled nuclei (blue, DAPI) and ECM proteins (Red, AF568) scale bar = 100μm. B) regions of human marrow with with fluorescently labelled nuclei (blue, DAPI) and ECM proteins (Red, AF568). Scale bar = 100 μm C) CD271 (green, AF488) and periostin (red, AF568) co-localisation in a bone-lining region with nuclei (blue, DAPI). individual channel images are shown. Dashed rectangle is shown as expanded view (bottom), scale bar = 50μm D) CD271 (green, AF488) and aggrecan (red, AF568) co-localisation in a bone lining region with nuclei (blue, DAPI), individual channel images are shown. Dashed rectangle is shown as expanded view (bottom), scale bar = 50μm. Bone regions marked as B.

We further investigated the distribution of periostin and aggrecan having identified these as the most differentially expressed proteins in Y201 secretome. We immunostained for CD271, one of the markers identified for appropriate selection and enrichment of colony forming human BMSCs that demonstrated tripotent differentiation *in vitro* [27, 28]. We identified CD271-positive staining in bone-lining regions with evidence for both aggrecan and periostin adjacent to the basal surfaces of these cells (Fig. 4 C&D).

### Extracellular matrix substrates of stromal populations regulates their migratory and morphological phenotype

To determine how differences in ECM composition identified in the LC-MS/MS analysis influenced ECM organisation, we examined the matrix substrate deposited by Y201 and Y202 BMSCs using scanning electron microscopy (SEM) and Focused Ion-Beam (FIB)-SEM. Y201 and Y202 BMSCs were cultured for 2 weeks to allow deposition of a layer of ECM onto the cell culture surface. After removal of the cell layer we used SEM to examine the topographical features of the matrices (Fig.5A). The matrix made by Y201 appeared to be more compact with larger and potentially deeper undulations. In contrast the matrix produced by Y202 cells appeared flattened, with fibres visible at both high and low magnifications (Fig. 5A). Differences in the organisation of Y201 and Y202 matrices was also demonstrated by FIB-SEM. The overall architecture of the ECM was apparent when observed at low magnification prior to initial FIB-SEM experiments. Y201 ECM appeared as a consistent mat of dense fibres whereas Y202 ECM presented as a more disperse meshwork with irregular patches of more fibrous matrix (Fig.5B). FIB-SEM was used to section through and image the ECM, revealing that ECM produced by Y201 cells was notably thicker than that produced by Y202 cells (Fig.5B).

**Figure 5.**
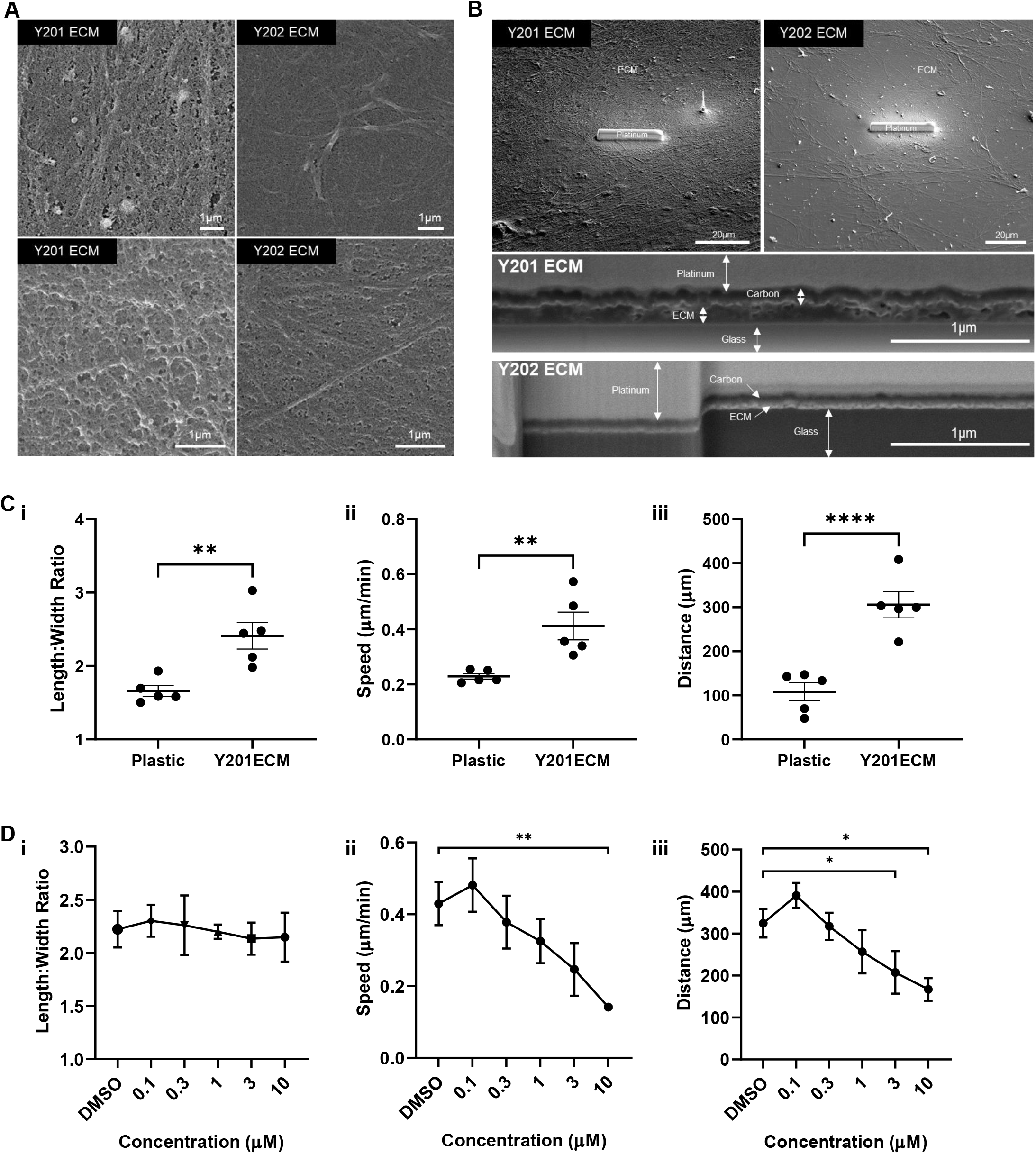
Effect of ECM substrates derived from Y201 BMSCs on Y202 BMSC migration. A) Scanning electron micrographs of Y201 and Y202 extracellular matrices after 2 weeks in culture. B) Scanning electron micrograph of expanded view of Y201 and Y202 extracellular matrices with platinum strip laid to protect sample during Focused Ion Beam (FIB) Milling. Bottom panels indicate side view after FIB milling revealing cross-section view of matrix deposition. C) Mean i) length:width ratio, ii) speed and iii) distance travelled of Y202 cells cultured on plastic or Y201 ECM (n=5 experiments). D) Mean i) length:width ratio ii) speed and iii) distance travelled of Y202 cells cultured on Y201 ECM with various concentrations of FAK inhibitor (PF573228) n=4 experiments. *p≤0.05, **p<0.01, ****p<0.0001. Error bars ± SEM

ECM deposited by BMSC subtypes appeared structurally distinct and so we hypothesised that this may explain the different migratory capacities of the producing cells. Indeed, Y202 cells seeded onto ECM from Y201 cells increased migration compared to those cultured on plastic. Y202 cells cultured on Y201 ECM became more fibroblastic as shown by the increased length:width ratio (Fig.5Ci). Y202 cells cultured on Y201 ECM also migrated further from their point of origin and at an increased speed versus Y202 cells grown on plastic (Fig.5Cii & iii). Next, we tested the role of focal adhesion kinase (FAK) in these ECM-mediated changes in migration. The FAK inhibitor (FAKi) (PF573228) did not significantly alter the length:width ratio of Y202 cells on Y201 ECM at any concentration tested (Fig.5Di) but did significantly reduce the mean migration speed (at 10μM FAKi) and the displacement of individual cells from their starting point Y202 cells treated (at 3 and 10μM FAKi) (Fig.5Dii-iii).

## Discussion

Our findings provide further evidence of a correlation between BMSC morphology and functionality, supporting previous evidence that morphologically distinct stromal subsets are likely to reflect functional heterogeneity, and observations that cells with different morphologies have altered inflammatory or differentiation characteristics [19, 29-32] We exploited a label-free ptychographic technique to track the morphology and motility of cells over time [33]. This could prove useful in the real-time discrimination of primary cell population phenotypes without the need for fluorescence-based or other end-point labelling methods. Using our simplified model of BMSC heterogeneity we showed that large, flat, inflammatory BMSCs were less motile than stem cell-like, spindle-shaped cells. In addition, cells matching these morphological parameters were reproducibly observed in primary cultures, suggesting that image-based morphometric analysis could be employed as a predictive measure of cell function, with previous evidence suggesting faster migrating BMSCs are indeed more likely to be multi-potent [34]. Furthermore, we demonstrated that the morphometric features of the atypical flattened BMSCs (Y202) were plastic and could be modified by exposure to factors secreted by more typical, spindle-shaped Y201 BMSCs. BMSC phenotypic plasticity may be, to some extent, determined by the secreted factors of the cell population as a whole, with ECM components being important determinants of cell behaviour.

The BMSC secretome is linked to cellular functionality, which is important both for the understanding of disease and potential uses of these cells in therapies [35]. We found that the secretome of multipotent Y201 BMSCs was strongly enriched for proteins involved in the production and modification of the ECM, as well as TGF-beta and Notch signalling pathways both of which are implicated in controlling BMSC differentiation [36, 37]. Subsequent assessment of the ECM produced by these BMSC lines identified a thicker and more complex matrix produced by Y201 cells, while Y202-derived matrices were relatively thin. The ECM has a prominent role in driving migration, and as such the increased production and secretion of matrix proteins captured in conditioned media could contribute to the phenotypic switch we saw in Y202 cells. Lumican, which was secreted at higher levels by Y202 cells, has previously been shown to inhibit the migration of MSCs, as well as regulation of immune responses in other cell types, potentially correlating with the slow moving immune-based role of Y202 [38].

Similarly, aggrecan and periostin which were more abundantly secreted in Y201 BMSCs compared to Y202 BMSCs, may act as candidate differentiators of cell phenotype. It is also possible that those proteins that did not differ significantly between the two BMSC lines represent a ‘core’ matrix common to all BMSC subtypes. The ECM is undoubtedly important for cellular function, mediating biochemical and mechanical signals to cells. Molecular patterning of a niche environment has previously been shown to regulate macrophages between a pro-healing and inflammatory phenotype [39]. This is likely to be of high importance for stem cells in a structurally diverse tissues such as bone marrow, where the role of ECM in maintaining hematopoietic stem cells in their niche has been increasingly characterised [40, 41]. Our identification of aggrecan and periostin underlying CD271^+^ cells in human bone marrow provides promising evidence that an *in vitro* matrix produced by cells isolated from a complex tissue may, at least in part, recapitulate the *in vivo* ECM composition, and may indeed contribute to a specialised stromal niche. Our findings are supported by previous evidence for CD271^+^ BMSC niches, with CD271^+^CD56^+^ cells found exclusively on trabecular bone surfaces, representative of an endosteal BMSC niche [42, 43]. CD271^+^ BMSCs are also 65-fold increased in BMSCs isolated from trabeculae versus bone marrow aspirates, again highlighting a more endosteal niche for this population [44]. The same pattern of CFU-F capacity and *in situ* localisation is seen when combined with another prospective potency marker, melanoma cell adhesion molecule (MCAM/CD146), as CD271^+^CD146^-/low^ populations were found as bone lining cells, whereas CD271^+^CD146^+^ were located perivascularly [45]. We hypothesise that differentiation-competent cells pattern their local environment with a phenotype-supportive matrix that is physically and biochemically suited to cell function, with our findings complementing other studies showing that matrix of young MSCs has been shown to restore proliferation and differentiation to older MSCs which has important implications for downstream therapeutic development [46-48].

Periostin has previously been linked with controlling the regenerative potential of periosteal skeletal stem cells, as well as supporting haematopoietic stem cells in the foetal liver niche and regulating their quiescence [49, 50]. The observation of rare periostin in bone marrow has not been previously reported in large-scale analyses of protein distribution across whole long-bones, however BMSC-derived periostin has also been shown in mouse to have functional effects in leukaemia, suggesting it is present in marrow [50-52]. Further, periostin knockdown in human BMSCs results in inhibition of osteogenic differentiation of these cells, indicating its importance for a differentiation-competent, stem cell phenotype.[53] The observation of periostin and aggrecan expression in trabecular bone regions in mouse and human tissue sections might also indicate conservation across species for these proteins in a stromal niche for bone lining cells. Follow up work to isolate CD271+ aggrecan and periostin-expressing primary BMSCs is necessary to determine if these possible biomarkers of potency are consistent and selective.

We also demonstrated that the ECM substrate produced by Y201 BMSCs may be partly responsible for its enhanced migratory phenotype; exposure of Y202 BMSCs to ECM derived from Y201 BMSCs resulted in increased migration, speed and distance in a FAK-dependent manner. The ability of the Y201 matrix to instruct the immotile Y202 to migrate highlights the importance of the secreted ECM in phenotypic plasticity and could correlate with previous observations of rejuvenation of cells by exposure to specific ECMs.[47]

In summary, we have demonstrated that there is a complex interplay between stromal cell subtypes that exhibit phenotypic plasticity driven by secreted signals, with the ECM playing a prominent role. As a result, the ECM will contribute to the initiation, maintenance and resolution of cellular heterogeneity. A stable and consistent ECM, for example at specific anatomical locations in vivo such as the endosteal niche, can also contribute to phenotypic stability.

## Experimental Procedures

### Cell Culture

Y201 and Y202 BMSC lines were cultured in complete medium (Dulbecco’s Modified Eagle Medium (DMEM) containing 10% foetal bovine serum (FBS), 100units/ml penicillin and 100μg/ml streptomycin) and incubated at 37°C in a 95% air/5% CO_2_ atmosphere. Cells were passaged using trypsin-EDTA on reaching 70-80% confluency. All work involving human samples was approved by the University of York, Department of Biology Ethics Committee. Primary human BMSCs were isolated from femoral heads obtained with informed consent during routine hip replacement or as explant cultures from the tibial plateau after routine knee replacement surgery.

### Conditioned Media Collection for secretome analysis and functional assays

Conditioned media was collected from 2x T175 flasks of Y201 and Y202 BMSC lines. Cells were grown to ∼80% confluency before washing 2x with PBS, 17ml of serum-free DMEM was added to the flasks and incubated as normal for 24h. Medium was collected and then centrifuged at 300g to remove any large cell debris. For functional assays, medium was stored at −80°C until required. For proteomic analyses, the medium was concentrated in 3kD-MWCO tubes (GE Healthcare) at 4500g until concentrated to ∼1ml in volume. Media were stored at −80°C until required.

### Preparation of MSC-derived Extracellular Matrix (ECM)

ECM was prepared from *in vitro* cell cultures using a protocol adapted from Ng, et al. [54]. Cells were seeded at 1000 cells/cm^2^ in 6-well plates or on 13mm glass coverslips and allowed to grow for 14 days. For days 1-7, cells were grown in complete medium and for days 8-14 this medium was supplemented with 50μM L-ascorbic acid (Sigma-Aldrich). Medium changes were performed every 3 days. On day 14, medium was aspirated and cells were removed from the deposited ECM by incubation (5 minutes, room temperature) with 20mM ammonium hydroxide with 0.5% Triton X-100 in PBS and gentle agitation every minute. Plates were washed 1x with PBS and 3x with sterile dH_2_O after cell clearing. Matrices were dried in a sterile laminar flow cabinet before storing at 4°C wrapped in parafilm for up to 1 month.

### Scanning Electron Microscopy (SEM)

ECM samples were fixed for 30 minutes in a mixture of 4% paraformaldehyde (PFA) + 2.5% glutaraldehyde in 100mM phosphate buffer (pH 7.0) at room temperature. Samples were washed twice for 10 minutes each with phosphate buffer before secondary fixation with 1% osmium tetroxide for 30 minutes at room temperature. Samples were washed twice with phosphate buffer for 10 minutes, then dehydrated in an ethanol series of 25%, 50%, 70%, 90% and 3 × 100% for 15 minutes at each stage. Samples were covered with hexamethyldisilazane for 15 minutes before aspirating and allowing to air dry. Samples were imaged with a JEOL 7800F Prime.

### Focused ion beam scanning electron microscopy (FIB-SEM)

Samples were prepared for FIB-SEM by fixing in 2.5% glutaraldehyde in 100mM phosphate buffer for 1h before 3 × 15 minute washes with phosphate buffer. A secondary fixation with 1% OsO_4_ in 100mM phosphate buffer was performed for 1h before 3 × 5 minute washes with ddH_2_O. Samples were then blocked in 1% uranyl acetate in ddH_2_O for 1h. Samples underwent dehydration in an ethanol series with 15 minutes in 30%, 50%, 70%, 90% and 2 × 15 minutes in absolute ethanol. The samples were then washed 2 × 5 minutes in epoxy propane before infiltrating with Epon-araldite resin (Epon 812, Araldite CY212) overnight. Excess resin was removed by spinning coverslips at 1000g before the resin was polymerised at 60°C for 48h. Prior to FIB milling, carbon coating was evaporated onto the matrix surface to provide a conductive sheath. The underlying film is protected from the destructive effect of the ion beam by the deposition of a thin (2 to 3 μm) layer of nanocrystalline platinum (Pt) and amorphous carbon. The Pt atoms provide a high-Z barrier to unwanted Ga ion exposure. Milling into the film commences with a high current ion probe (7 nA) that produces a deep, triangular trench to a depth of several micrometres. A series of ‘cleaning scans’ were executed with smaller ion probe currents (1 nA, 300 pA, 50 pA, all at 30 keV) to remove thin layers of damaged surface material. This exposed the interfaces between the substrate, the thin film and the deposited carbon and Pt layers. Finally, the sample could be tilted to ensure that optics were as close to the milled surface as possible for imaging.

### Proteomic Analysis

Concentrated whole secretome samples were added to 8M urea with 20mM HEPES, 1mM sodium orthovanadate, 1mM β-glycerophosphate and 2.5mM sodium pyrophosphate (Sigma-Aldrich). Protein was in-solution reduced and alkylated before digestion with a combination of Lys-C and trypsin proteases. Resulting peptides were analysed over 1 h LC-MS acquisitions using an Orbitrap Fusion. Peptides were eluted into the mass spectrometer from a 50 cm C18 EN PepMap column. Three biological replicates for each cell line were run. Tandem mass spectra were searched against the human subset of the UniProt database using Mascot and peptide identifications were filtered through the Percolator algorithm to achieve a global 1% false discovery rate (FDR). Identifications were imported back into Progenesis QI and mapped onto MS1 peak areas. Peak areas were normalised to total ion intensity for all identified peptides. Relative protein quantification was performed using relative peak areas of non-conflicting peptides. Relative fold differences and associated p-values for differential abundance were calculated in Progenesis QI.

### Bioinformatic analyses

Proteins were annotated for involvement in the Matrisome using the MatrisomeDB database at www.http://matrisomeproject.mit.edu/ [55]. Chi-squared tests were performed in Graphpad Prism.

Lists of significantly more abundant genes and proteins were analysed for pathway enrichment against the curated Kyoto Encyclopedia of Genes and Genomes (KEGG) database using the Molecular Signatures Database website on version 7.2 [56-58]. Enrichment was performed for significantly different protein lists and results filtered to exclude terms with FDR corrected p-values (q) of >0.05. To minimise the effect of confounding and relatively uninformative terms, a filter excluded protein-sets containing more than 500 proteins. Where p-values for enriched pathways were the same, samples were ordered by the MSigDB k/K ratio where k = the number of proteins identified in the protein-set and K = the total number of proteins in that set. Enrichments were presented in bar-charts generated in Graphpad Prism. Cytoscape was used for visualisation of cellular location of proteins from secretomics [59].

### Ptychography, cell tracking and image analysis

For cell migration and morphology analysis cells were seeded as 6-well colony-forming unit fibroblastic (CFU-F) assays and Ptychography was performed using a PhaseFocus VL21 Livecyte imaging platform for live cell tracking analysis. Images were taken at 20-26-minute intervals for 96 hours. Images were first processed with a rolling ball algorithm before smoothing was applied to remove low frequency noise. Points of maximal brightness, indicating areas of high phase-contrast corresponding to cell nuclei, were identified in the smoothed image and were used as seeding points for the identification of individual cells. Seed points were consolidated where points that did not change in pixel intensity within a threshold were removed, this enabled removal of multiple seed points in a single cell. Thresholding and segmentation levels were set to define the cell area against the background. This processing pipeline was applied to all images in an experiment. The output images then allowed tracking of cells and using a spatial and temporal dot plot, along with quantification of various morphological metrics such as dry-mass, area, width and length. Small debris was removed by an exclusion gate removing objects that were less than 250pg in dry mass and less than 1000μm^2^. Large doublets and debris were excluded with an area over 25,000μm^2^. Manual removal of debris was also performed by visual assessment. To be included in analyses, cells had to be tracked for a minimum of 20-frames. Cell morphology and migration was quantified using the PhaseFocus analysis platform and statistical tests performed in Graphpad Prism. Rose plots were generated using the mTrackJ plugin in ImageJ [60]. The image analysis programme CellProfiler was used to generate a pipeline to assess the morphological characteristics of BMSCs [61]. This pipeline was subsequently used to categorise different BMSC subtypes into subgroups of Y201, Y202 or a group of cells that were between categories.

### CFU-F assays and image analysis

For CFU-F assays, cells were seeded at 10 cells/cm^2^ in 6-well plates using DMEM supplemented with 20% Hyclone FBS containing 100units/ml penicillin, 100μg/ml streptomycin. Conditioned medium for use in the CFU-F assays was collected from Y201 and Y202 MSCs by incubating in serum-free medium at ∼80% confluency for 24 hours before collecting media, centrifuging at 300g to remove cell debris, and counting the number of cells. The conditioned medium was then diluted with additional serum-free DMEM to give 12ml conditioned media/million cells. This medium was then supplemented with a final concentration of 20% Hyclone FBS for use in CFU-F assays. For CFU-Fs, primary cells and cell lines were seeded in unconditioned Hyclone medium before media changes were performed every 4 days post-seeding and plates were fixed and stained at day 10 for cell lines and day 14 for primary cell. Plates were stained with (0.05% crystal violet + 1% formaldehyde + 1% methanol in PBS) for imaging or were washed 1x with PBS and the cells lysed with 350μL of RA1 cell lysis buffer (Machery-Nagel) + 3.5μL β-mercaptoethanol for every 3 wells. Well plates were air dried before scanning on an Epson Perfection 4990 Photo scanner at 1200dpi. A CellProfiler pipeline was subsequently developed to detect and measure colonies accurately. The scanned image was loaded into CellProfiler and converted to a greyscale image using the ColorToGrey module, splitting the image into Red, Green and Blue channels. The Blue channel was then thresholded to 0.99 to include all features identified as completely black. Well edges were identified as primary objects of size 1000-2000-pixel units in diameter and with a manual threshold of 0.99 to include all features, this reproducibly identified the well edges as primary objects. In order to fit this as a complete circle a grid was defined using DefineGrid and then true circles were placed using the IdentifyObjectsinGrid module. The circle was shrunk by 10 pixels in diameter to prevent running over the edge of the well. The UnmixColors module was used to create an image without any Blue absorbance (Red and Green absorbance of 1, Blue absorbance of 0). The area of this image outside of the wells was cropped using the 10 pixel shrunken circles. Illumination correction was calculated (block size 20, median filter and Object size filter with median object size of 80 pixels), and applied by subtraction. The edges of features were enhanced using the Sobel method in the EnhanceEdges module which identified cells that had dispersed away from an otherwise tight colony. The distance of these cells was then closed using a Closing module in a Diamond shape with a reach of 10 pixels. Colonies were subsequently detected by an IdentifyPrimaryObjects module with typical diameter between 60-800 pixels and using the RobustBackground with a Mode averaging. Manual correction of colony detection could then be applied in CellProfiler. Resultant colonies were measured for size and shape characteristics and used as a mask to analyse other features of the colonies such as intensity.

### Focal adhesion immunostaining assessment

Y201 and Y202 cells were plated onto glass coverslips left to adhere for 24h. Cells were fixed briefly in 4% methanol-free PFA in PBS before washing 3 x with PBS. Cells were permeabilised in 0.1% Triton X-100 in PBS for 30 minutes and washed 3 x with PBS. Cells were then blocked for 30 minutes with 10% goat serum in PBS. Anti-vinculin antibody (Sigma-Aldrich) was added in 1% BSA and incubated at room temperature for 1 hour. Cells were washed 3x with PBS before Goat anti-mouse Alexafluor-488 conjugated secondary antibody (Thermofisher) was added along with Cruzfluor-594 conjugated phalloidin (Santa Cruz) for 45 minutes in PBS followed by another 3x washes. Nuclei were counterstained with 0.2μg/ml DAPI for 10 minutes before rinsing briefly in distilled water and leaving to air-dry. Coverslips were mounted onto a microscope slide with Prolong gold antifade (ThermoFisher).

Slides were imaged on a Zeiss LSM880 or LSM780 microscope. Focal adhesion sizes were quantified using ImageJ. All antibody manufacturers, clones and dilutions can be found in supplementary Table S1.

### Immunofluorescence of mouse femurs

Femurs from 8-12 week old C57BL/6 mice were collected and fixed in 4% PFA in PBS for 24 hours followed by decalcification in 10% EDTA in PBS pH 7.5 for 24 hours. Bones were then transferred to 30% sucrose in PBS for 24 hours before freezing in optimal cutting temperature (OCT) compound on a dry ice and ethanol slurry. Sections were cut to 8μm thickness on a Bright OTF5000 cryostat and collected on Superfrost plus slides (ThermoFisher). Sections were blocked in 10% goat serum + 0.1% Tween-20 in PBS for 45 minutes before addition of primary antibodies in 1% Bovine Serum Albumin + 0.05% Tween-20 and left overnight at 4°C. Sections were washed 3 times for five minutes with PBS before adding all secondary antibodies in PBS for 1 hour at room temperature. Antibody manufacturer and dilution details are provided in supplementary Table S1. Three five-minute washes were performed before staining for 10 minutes with 0.2μg/ml DAPI in PBS (Sigma). Slides were rinsed in dH_2_O and dried before mounting a glass coverslip with Prolong Gold antifade mounting medium (Invitrogen). Images were taken on LSM880 or LSM780 confocal microscopes or a Zeiss AxioScan slidescanner (Zeiss).

### Immunofluorescence of human bone

Human femoral heads from routine hip and knee replacements were donated following informed consent from Clifton Park Hospital under ethical approval from the local NHS Research Ethics Committee and the University of York, Department of Biology Ethics Committee. A CleanCut bone saw (deSoutter medical) was used to cut femoral heads which were then dissected into roughly 1cm^3^ pieces using a fresh scalpel. Processing steps were carried out at 4°C. Bone pieces were fixed in 4% PFA for 24 hours. After fixation, bone pieces were washed once with PBS before decalcifying for 48 hours in 10% EDTA in PBS at pH 7.5. Bone pieces were then cryoprotected by submerging in 30% sucrose in PBS for 24 hours. Bone pieces were cut into smaller pieces with a scalpel before embedding in OCT on a dry ice ethanol slurry and cutting in a Bright OTF5000 cryostat. Sections were cut at 10μm thickness and collected onto Superfrost plus slides (Thermofisher). Immunofluorescent staining was then performed as for the sections of mouse bone described above. All antibody manufacturers, clones and dilutions can be found in supplementary Table S1.

## Acknowledgements

This work was funded by the Biotechnology and Biological Sciences Research Council (BBSRC) United Kingdom, Doctoral Training Partnership grant (BB/M011151/1) and the Tissue Engineering and Regenerative Therapies Centre Versus Arthritis (21156). We thank staff at University of York Technology Facility for their help in performing the LC-MS/MS and for the use of the PhaseFocus Livecyte and confocal microscopes. We would also like to thank Adam Dowle of The The York Centre of Excellence in Mass Spectrometry which was created thanks to a major capital investment through Science City York, supported by Yorkshire Forward with funds from the Northern Way Initiative, and subsequent support from EPSRC (EP/K039660/1; EP/M028127/1). We also thank Jon Barnard for his assistance with FIBSEM at the York Nanocentre and Richard Kasprowicz for writing the code used to make the rose plots. We are also very grateful to the staff and patients from Clifton Park Hospital York for providing primary tissue from which BMSCs were extracted.

## Contributions

**Andrew Stone:** Conceptualization; Data curation; Formal analysis; Investigation; Methodology; Validation; Visualization; Writing - original draft; Writing - review & editing. **Emma Rand:** Data curation; Formal analysis; Investigation; Methodology; Visualization, Writing - review & editing. **Gabe Thornes:** Data curation; Formal analysis; Investigation; Methodology; Writing - review & editing. **Alasdair Kay:** Data curation; Formal analysis; Investigation; Methodology; Writing - review & editing. **Amanda Barnes:** Investigation; Methodology; Writing - review & editing. **Ian Hitchcock:** Conceptualization; Funding acquisition; Methodology; Resources; Software; Supervision; Writing - review & editing. **Paul Genever:** Conceptualization; Formal analysis; Funding acquisition; Methodology; Project administration; Resources; Supervision; Writing - original draft; Writing - review & editing.

---

**Supplementary Figure 1.**
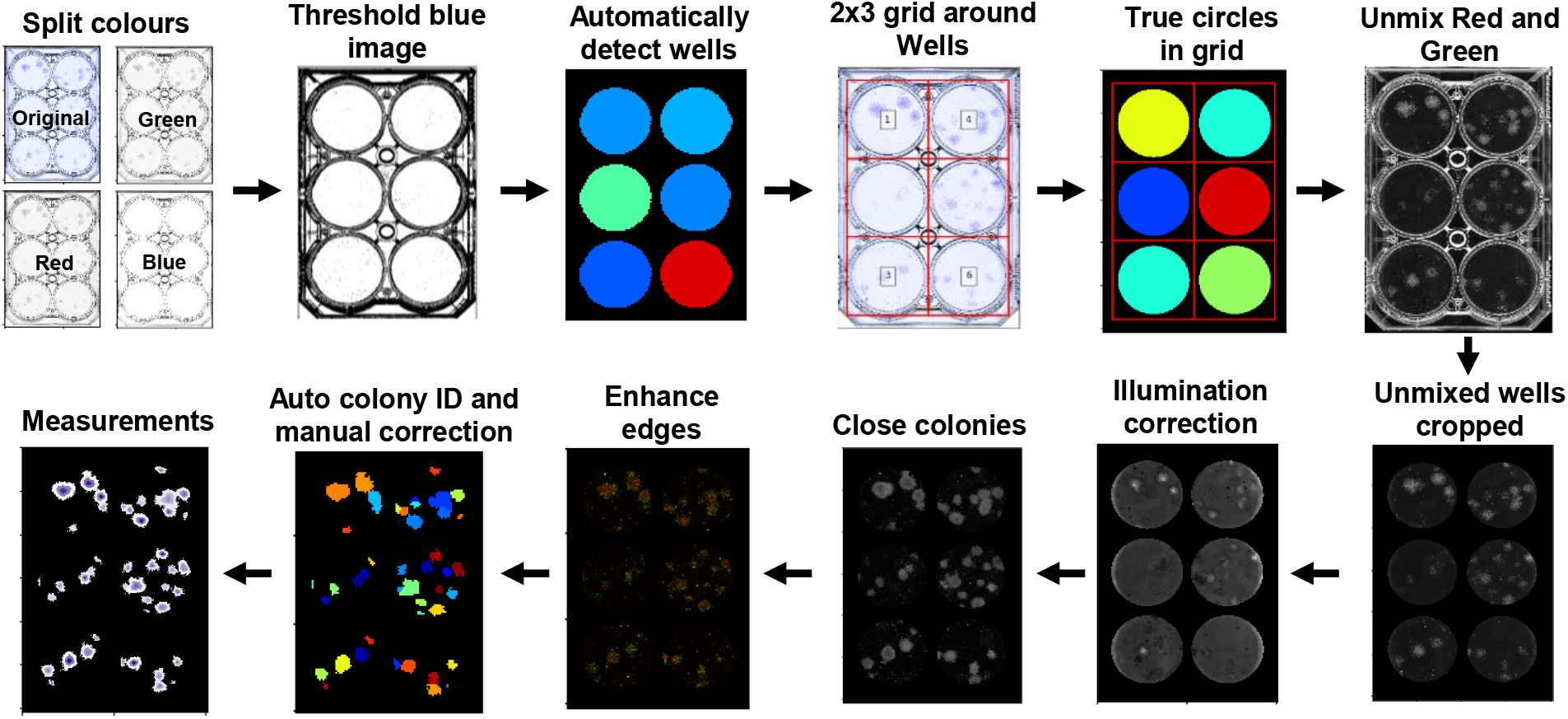
Outline of the CellProfiler image analysis pipeline developed for semi-automated detection and quantification of crystal violet stained CFU-F assays in 6 well plates.

**Supplementary Figure 2.**
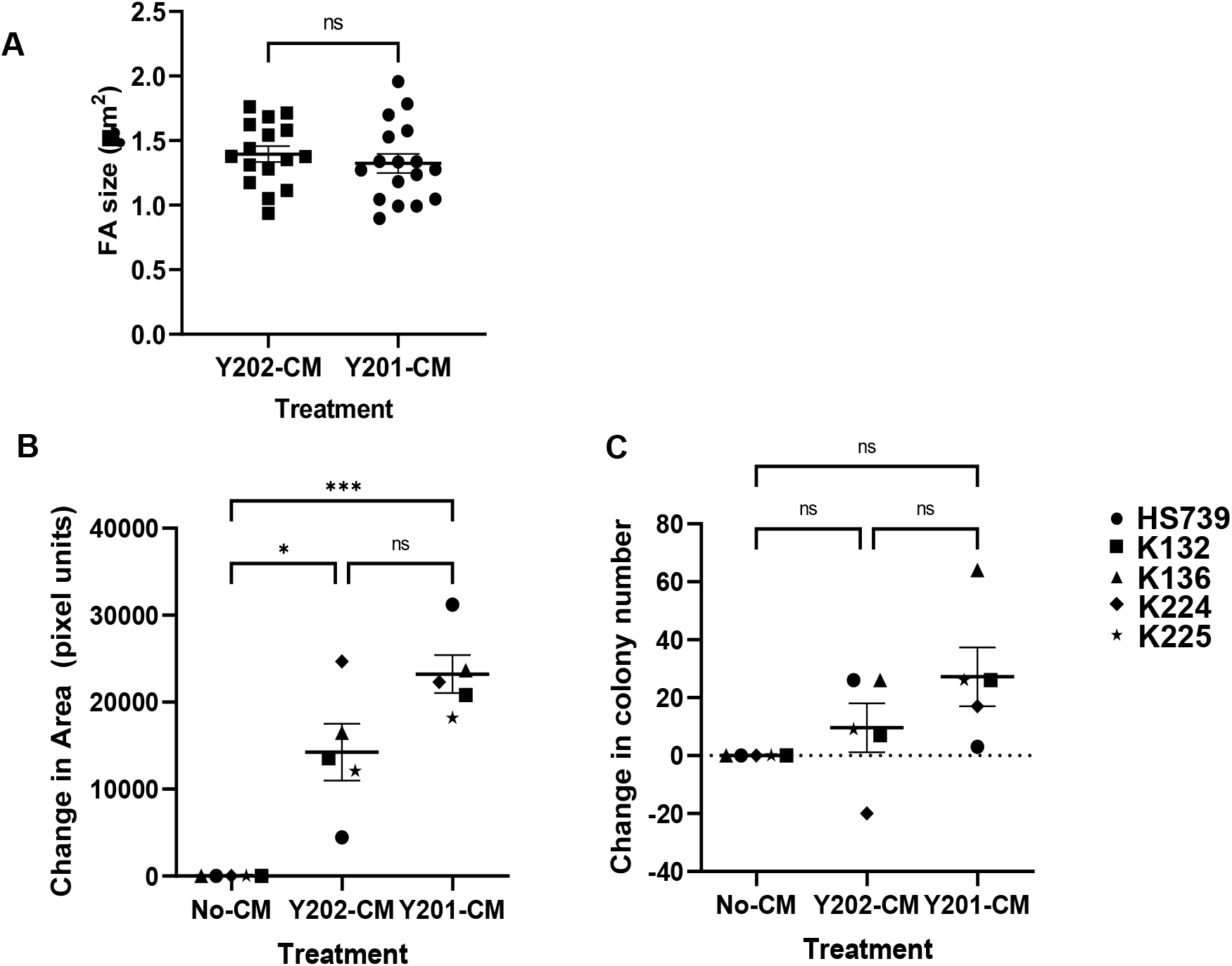
A) Quantification of focal adhesion size following fluorescent staining of Y202 cells treated with either Y201-CM or Y202-CM for 24 hours. B) Mean colony area from *in vitro* aged primary BMSC CFU-Fs (n=5) (ANOVA: *F=3*.*863, df = 1*.*833, 7*.*332 p = 0*.*0738*) C) Total colonies identified from CFU-Fs of *in vitro* aged primary BMSCs (Friedman test, *p* = 0.0085) with post-hoc test revealing significant effect of Y201CM vs no CM (*p = 0*.*0044)* n = 5. *p≤0.05, ***p<0.001, ns = not significant. Error bars ± SEM.

**Supplementary Figure 3.**
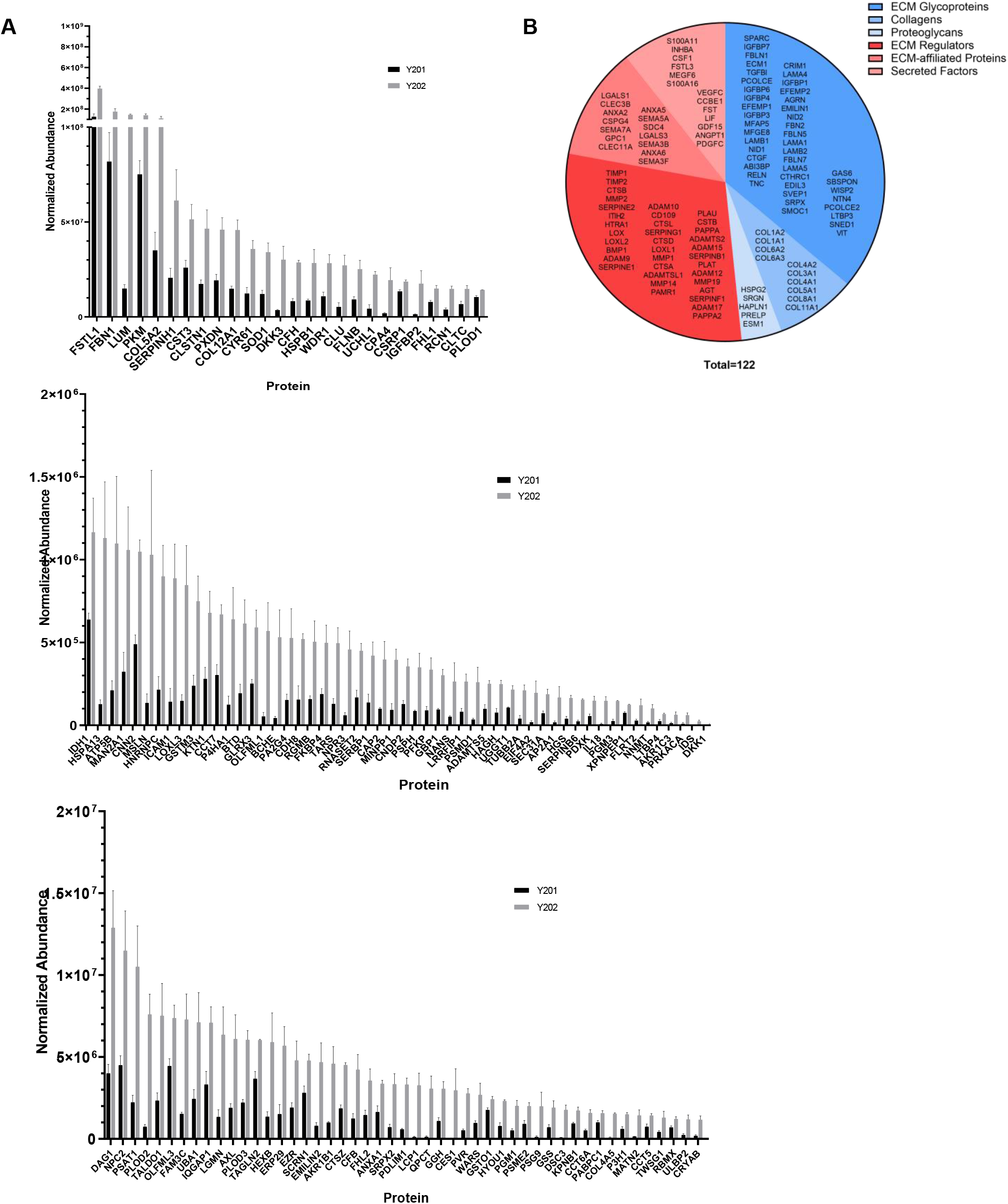
A) Significantly enriched proteins secreted by Y202 versus Y201 represented in order of normalised abundance from LC-MS/MS. Graphs are split for ease of interpretation while maintaining a linear scale, Means ± SEM. B) Matrisome annotated proteins that were not significantly altered between Y201 and Y202 secretomes. Proteins are labelled as core-matrisome (blue) or matrisome-associated (red) with shading representing sub-categories of these annotations.

**Supplementary figure 4.**
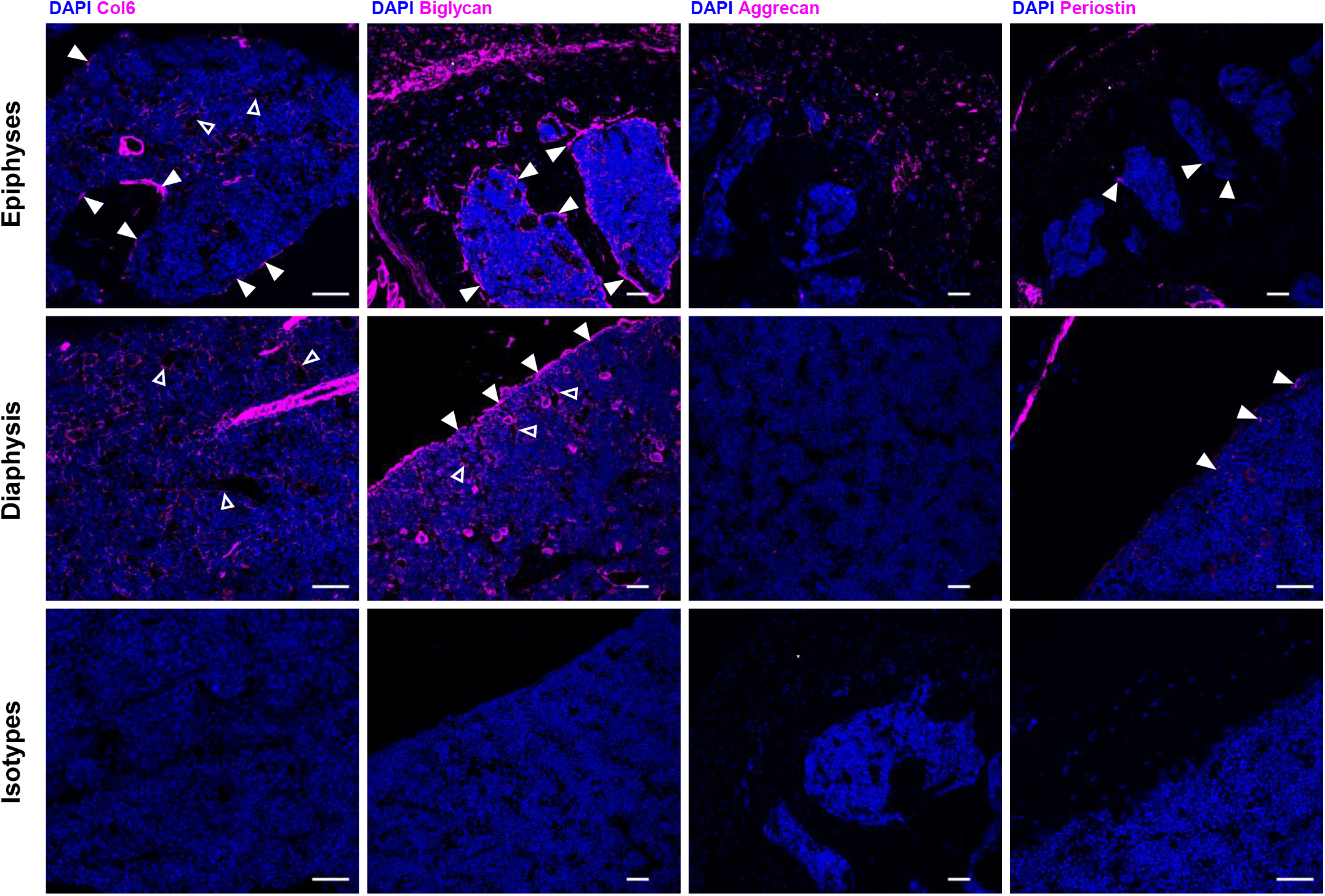
Representative immunofluorescence microscopy images of various ECM proteins (violet) in mouse bone marrow with DAPI (blue) nuclear stain. Top row shows imaging of epiphyseal region of mouse femur. Middle row shows regions from the diaphysis of the femur. The bottom row shows isotype controls for respective stains. Expression along the endosteal surface is marked by closed arrowheads. Expression of protein around possible endothelial vessels is marked by open arrowheads. Scale bars = 50μm.

