## Supplementary figures and images for "Extracellular matrices of bone marrow stroma regulate cell phenotype and contribute to distinct stromal niches *in vivo*"

### Supplemental Video 1

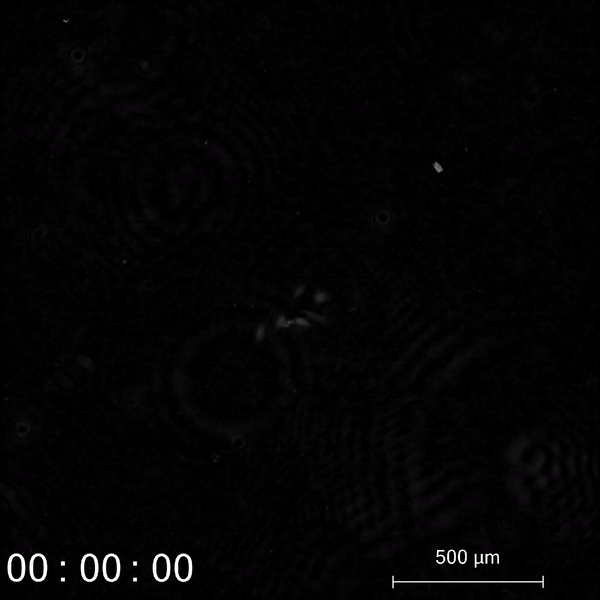

### Supplemental Video 2

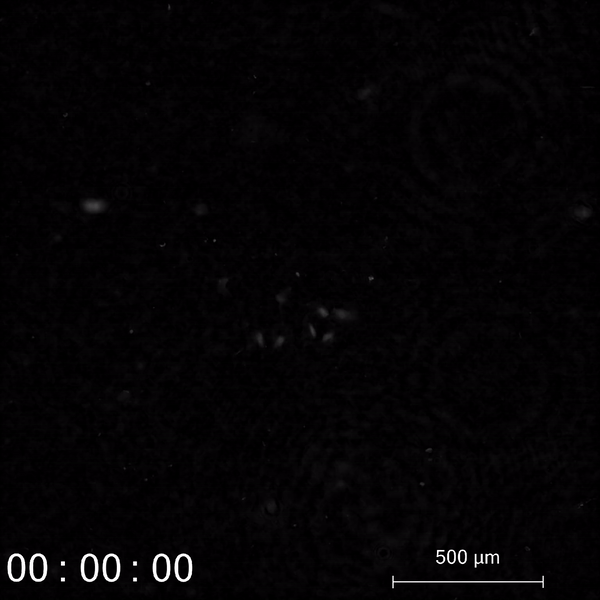
